# Engineering highly multivalent sperm-binding IgG antibodies for potent non-hormonal female contraception

**DOI:** 10.1101/2020.04.22.055301

**Authors:** Bhawana Shrestha, Alison Schaefer, Jamal Saada, Zhu Yong, Timothy M. Jacobs, Elizabeth C. Chavez, Stuart S. Omsted, Kathleen Vincent, Thomas R. Moench, Samuel K. Lai

## Abstract

Many women risk unintended pregnancy due to dissatisfaction with available hormonal contraceptive methods. This led us to pursue topical sperm-binding monoclonal antibodies as a strategy for safe, non-hormonal contraception. Motivated by the greater agglutination potencies of polymeric immunoglobulins such as IgM and the exceptional bioprocessing ease in manufacturing IgG, we engineered IgGs possessing 6-10 Fabs against a unique surface antigen universally present on human sperm. These highly multivalent IgGs (HM-IgGs) are at least 10- to 16-fold more potent and faster than the parent IgG at agglutinating sperm, while preserving Fc-mediated trapping of individual spermatozoa in mucus. The increased potencies translate to effective (>99.9%) reduction of progressively motile sperm in the sheep vagina using 33 micrograms of the 10 Fab HM-IgG. HM-IgGs produce at comparable yields and possess identical thermal stability to the parent IgG, with greater homogeneity. HM-IgGs represent not only promising biologics for non-hormonal contraception but also a promising platform for generating potent agglutinating mAb for diverse medical applications.

## Introduction

Nearly half of all pregnancies in the United States are unintended despite the availability of cheap and effective hormonal contraceptives, which creates an enormous cost burden on the healthcare system^1,2^. Many women avoid hormonal contraception due to real and perceived side effects including increased risks of breast cancer, depression, prolonged menstrual cycle, nausea and migraines^3,4^. Many women also have medical contraindications to the use of estrogen-based hormonal contraceptives^5–7^. Thus, there is a strong unmet need for alternative, non-hormonal contraceptives.

An effective non-hormonal contraceptive mechanism already exists in nature: anti-sperm antibodies (ASAs) in the female reproductive tract (FRT) of infertile women can trap vigorously motile sperm in mucus and prevent them from reaching the egg, via two distinct mechanism^8,9^. First, at high sperm concentration, ASA can agglutinate sperm into clusters that are too large to penetrate mucus, a process particularly potent with polymeric antibodies (Abs) such as IgM^10–12^. Second, at lower sperm concentration, ASA can trap individual spermatozoa in mucus by forming multiple low affinity Fc-mucin bonds between sperm-bound ASA and mucin fibers^13–15^. Years ago, these observations motivated the development of contraceptive vaccines^16–19^. Vaccines eliciting ASA offered considerable contraceptive efficacy, but the approach stalled due to unresolved variability in the intensity and duration of the vaccine responses in humans, as well as concerns that active vaccination might lead to irreversible infertility^20,21^. In contrast, sustained delivery of ASA at pharmacologically active doses locally in the vagina can overcome many of the key drawbacks of contraceptive vaccines, making possible both consistently effective contraception and rapid reversibility. Indeed, vaginal delivery of sperm agglutinating IgM in the highly fertile rabbit model reduced embryo formation by 95%^22^.

Unfortunately, this topical passive immunocontraception approach has never been tried in humans, due in part to manufacturing and stability challenges with IgM and the limited potencies with IgG Abs. We hypothesized that we can engineer a stable and potently agglutinating ASA for non-hormonal contraception by linking multiple Fabs to a parent IgG, creating highly multivalent IgGs (HM-IgGs) with precisely tunable valencies^23^. Here, we report the development of three potent and stable highly multivalent anti-sperm IgGs: Fab-IgG-Fab (FIF;6-Fabs), Fab-IgG-Fab-Fab (FIFF; 8-Fabs) and Fab-Fab-IgG-Fab-Fab (FFIFF; 10-Fabs) **(Fig. 1a**). These constructs all exhibit superior (>10-fold) agglutination potency than IgG and effectively reduce progressively motile (PM) sperm in the sheep vaginal model.

**Fig. 1:**
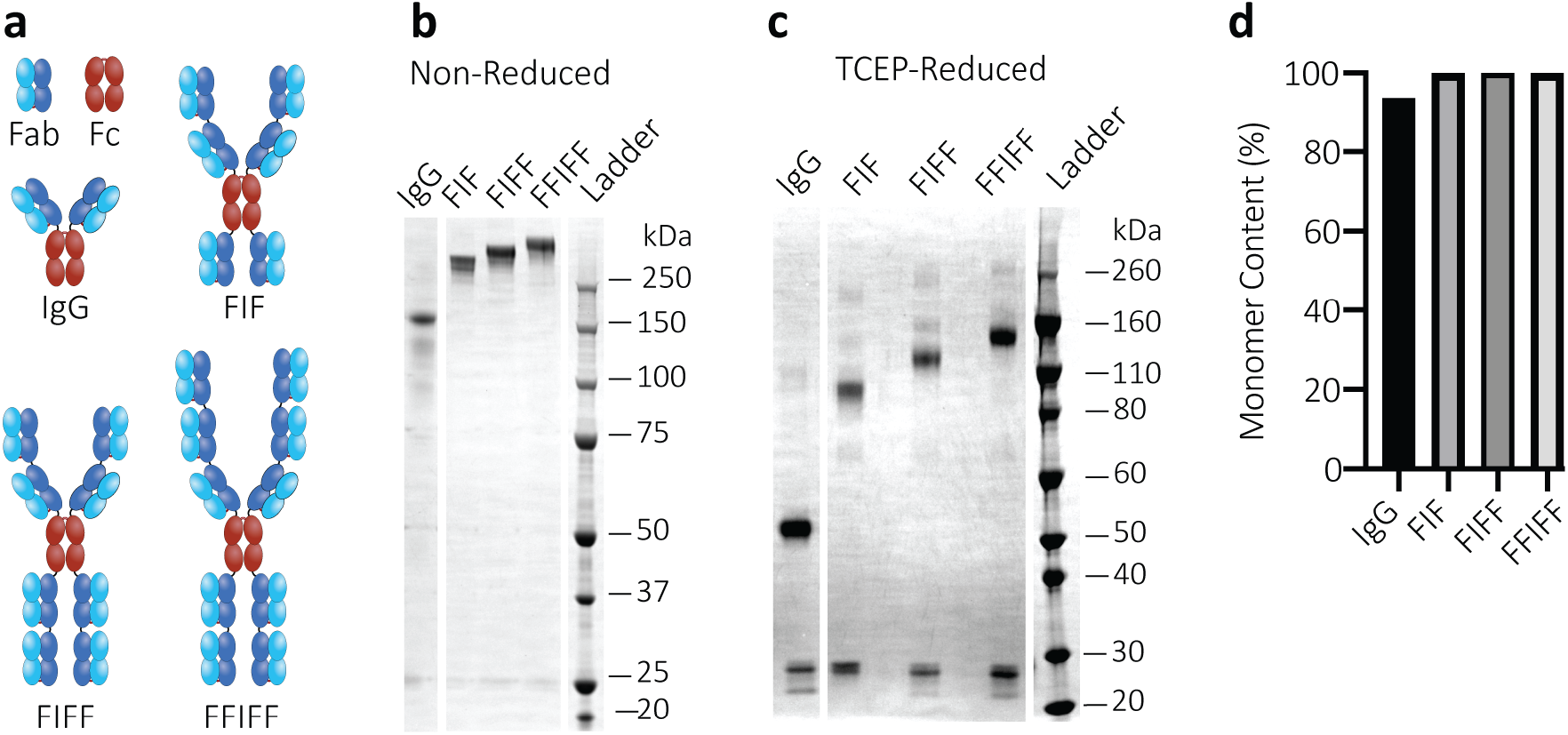
Production and characterization of highly multivalent anti-sperm IgG antibodies. (a) Schematic diagrams of anti-sperm IgG, Fab-IgG-Fab (FIF), Fab-IgG-Fab-Fab (FIFF) and Fab-Fab-IgG-Fab-Fab (FFIFF). In HM-IgGs, the additional Fab/s are linked to the N-terminal and C-terminal of baseline IgG using flexible glycine-serine linkers. (b) Non-reducing and (c) Reducing SDS-PAGE analysis of the indicated Abs (1 μg) after expression in Expi293 cells and purification by protein A/G chromatography. Non-reducing SDS-PAGE showcases the total molecular weight of the Abs. Tris (2-carboxyethyl) phosphine hydrochloride (TCEP)-mediated reducing SDS-PAGE displays the molecular weight of individual heavy chain and light chain of Abs. The experiment was repeated independently two times with similar results. (d) Demonstration of the purity and homogeneity of the indicated anti-sperm Abs (50-100 μg) using Size Exclusion Chromatography with Multiple Angle Light Scattering (SEC-MALS) analysis. Y-axis indicates the total percentage of Abs representing their theoretical molecular weights. Data collected from one experiment.

## Results

### Generation of FIF, FIFF and FFIFF mAbs

Human sperm are coated with a unique glycoform of CD52^24^ (hereafter referred to as CD52g) that is produced and secreted by epithelial cells lining the lumen of the epididymis, and present only on sperm, semen white blood cells, and the epithelium of the vas deferens and seminal vesicles^25^. The CD52g antigen appears to be universally present on all human sperm while absent in all other tissues, and absent in women^25^. We engineered different HM-IgG molecules with a Fab domain previously isolated from a healthy but immune infertile woman^10,26^; this Fab rapidly bound sperm in 100 out of 100 semen specimens **(Supplementary Table 2)**, with Fab domains interspersed by flexible glycine-serine linkers. Anti-CD52g FIF, FIFF and FFIFF all expressed at levels similar, if not exceeding those of the parent IgG when produced via transient expression in Expi293F cells **(Supplementary Fig. 1a)**. All 3 constructs properly assembled at their theoretical molecular weights **(Fig. 1b & 1c**) and were highly homogenous **(Fig. 1d & Supplementary Fig. 1b)**, with even greater homogeneity than the parent IgG, which possesses a small fraction of aggregates. All 3 constructs were highly stable, with comparable melting temperatures (T_m_) and aggregating temperatures (T_agg_) to the parent IgG. All mAbs unfolded (denatured) at similar high temperatures of Tm_1_ ≥ 71.1°C and Tm_2_ ≥ 80°C, while aggregation began only at ≥ 80°C (**Supplementary Fig. 1c**), with slightly increased T_m_ and T_agg_ for the 3 HM-IgG constructs. All constructs bound to human sperm in a whole-sperm ELISA assay, with apparent greater avidity of the FIFF and FFIFF multivalent constructs than the parent IgG at lower mAb concentration (**Supplementary Fig. 1d**).

### HM-IgGs exhibit greater agglutination potency than IgG

Progressively motility is crucial for sperm, both to enable sperm to swim through mucus to reach the egg as well as to penetrate the zona pellucida to fertilize the egg. We first assessed the agglutination potencies of all mAbs using a sperm escape assay, which quantifies, using Computer Assister Sperm Analysis (CASA), the number of PM sperm that escaped agglutination over 5 mins when mixed with specific mAbs or sperm washing media control^27,28^. At the final concentration of 5 million PM sperm/mL reflecting typical concentrations of PM sperm in fertile males^29,30^, all 3 HM-IgG constructs exhibited at least 16-fold greater agglutination potency than IgG, defined as the minimal mAb concentrations at which PM sperm are reduced by >98%. The minimum concentration of IgG needed was ~6.25 μg/mL, whereas all 3 HM-IgG constructs were able to do so down to 0.39 μg/mL **(Fig. 2a).** The greater agglutination potency of HM-IgG constructs was visually confirmed using scanning electron microscopy (SEM) on the mAb-treated washed sperm **(Supplementary Fig. 2).** To ensure efficient agglutination occurs not just with the washed sperm but also with native semen, we further assessed the agglutination of the most potent construct (FFIFF) vs IgG. Both FFIFF and the parent IgG required higher mAb concentration to reduce PM sperm quantities by >98% in whole semen, likely due to the greater amounts of CD52g present on both non-PM sperm as well as on exosomes in the seminal plasma. Nevertheless, FFIFF remained approximately 16-fold more potent than IgG **(Fig. 2c)**.

**Fig. 2:**
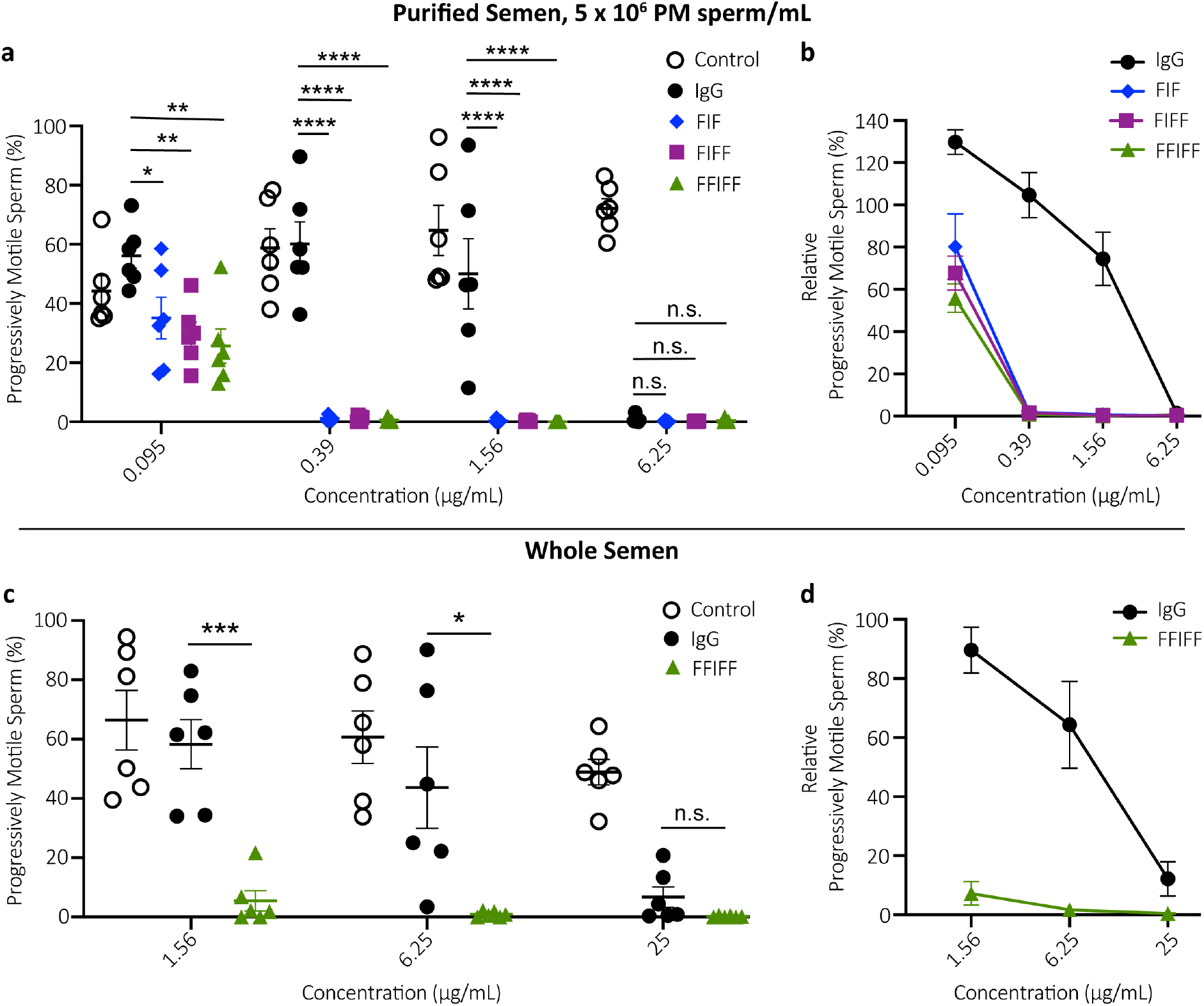
Multimerization markedly enhances the agglutination potency of anti-sperm IgG antibodies. (a) Sperm agglutination potency of the parent IgG, FIF, FIFF and FFIFF measured by the CASA-based quantification of the percentage of sperm that remains progressively motile (PM) after Ab-treatment compared to pre-treatment condition. Purified motile sperm at the concentration of 5 x 10^6^ PM sperm/mL was used. (b) The sperm agglutination potency of the Abs against 5 x 10^6^ PM sperm/mL normalized to the sperm washing media control. (c) Further assessment of sperm agglutination potency of the parent IgG and FFIFF using whole semen. (d) The sperm agglutination potency of the parent IgG and FFIFF against whole semen normalized to the sperm washing media control. Data were obtained from n = 6 independent experiments with at least n = 4 unique semen donors. Each experiment was performed in duplicates and averaged. P values were calculated using a one-way ANOVA with Dunnett’s multiple comparisons test. *P < 0.05, **P < 0.01, ***P < 0.001 and ****P < 0.0001. Lines indicate arithmetic mean values and standard error of mean.

### HM-IgGs induce faster agglutination kinetics than IgG

For effective vaginal immunocontraception, mAbs must agglutinate/immobilize sperm before they reach the upper reproductive tract; thus, the speed with which PM sperm become agglutinated will likely correlate with contraceptive efficacy^31^. We therefore quantified the kinetics of sperm agglutination immediately following mixing of sperm and mAb using CASA with washed sperm at a standard concentration of 5 million PM sperm/mL. The parent IgG reduced PM sperm by ≥90% within 90 s in 5 of 6 semen samples at 6.25 μg/mL but failed to do so in 6 of 6 samples at 1.56 μg/mL **(Fig. 3a).**

**Fig. 3:**
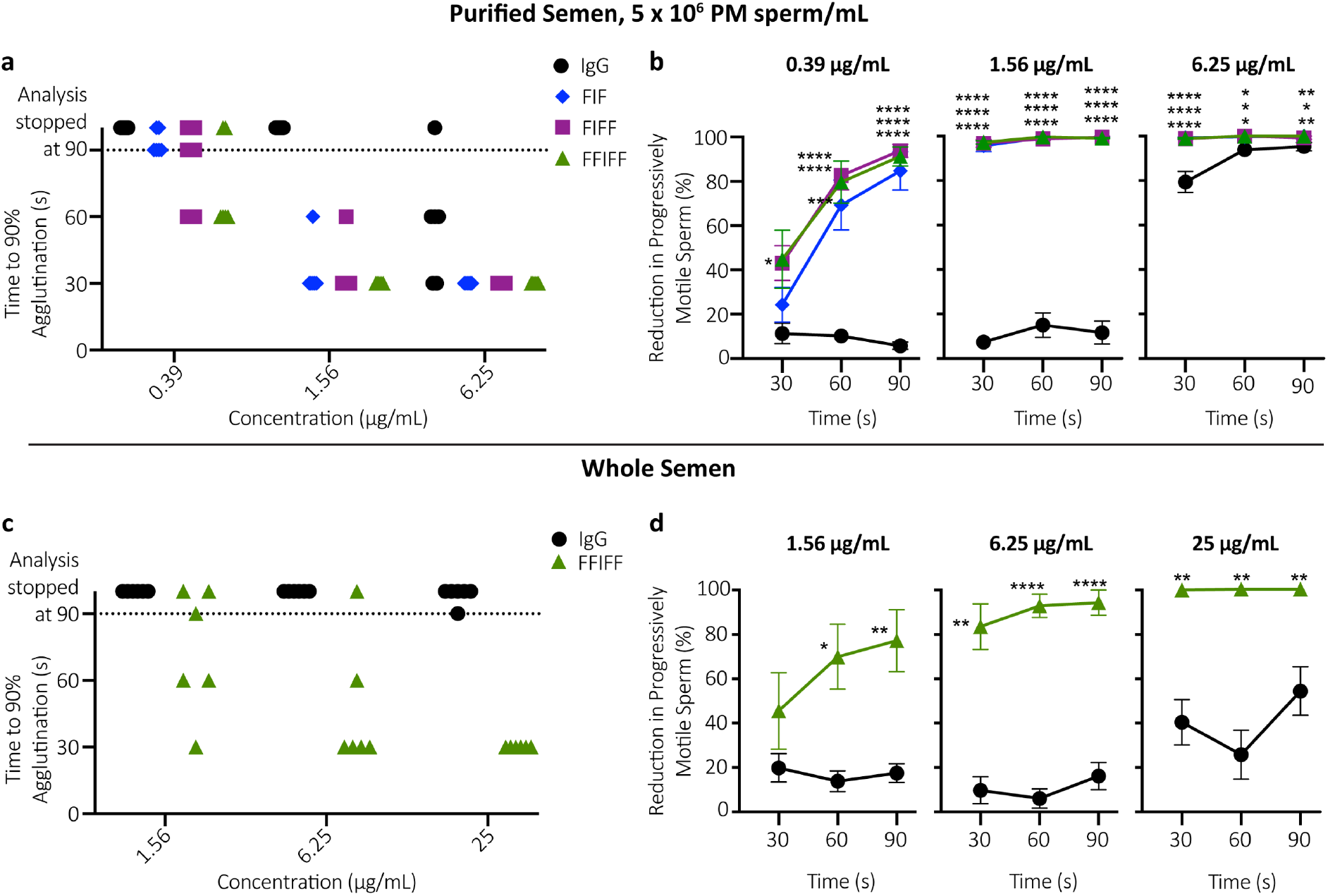
Multimerization significantly accelerates the agglutination kinetics of anti-sperm IgG antibodies. (a) Sperm agglutination kinetics of the parent IgG, FIF, FIFF and FFIFF measured by the quantification of time required to achieve 90% agglutination of PM sperm compared to sperm washing media control using 5 x 10^6^ PM sperm/mL. (b) The rate of sperm agglutination for IgG, FIF, FIFF and FFIFF determined by the reduction in percentage of PM sperm at three different timepoints after Ab-treatment compared to sperm washing media control using 5 x 10^6^ PM sperm/mL. (c) Sperm agglutination kinetics and (d) The rate of sperm agglutination for IgG and FFIFF evaluated using whole semen. Data were obtained from n = 6 independent experiments with at least n = 4 unique semen donors. Each experiment was performed in duplicates and averaged. P values were calculated using a one-way ANOVA with Dunnett’s multiple comparisons test for Fig. 3b and a one-tailed t-test for Fig. 3d. *P < 0.05, **P < 0.01, ***P < 0.001 and ****P < 0.0001. Lines indicate arithmetic mean values and standard error of mean.

In contrast, all 3 HM-IgG constructs agglutinated ≥90% of PM sperm within 60 s in all cases at both 1.56 μg/mL and 6.25 μg/mL concentrations, and only began to fail to do so at 0.39 μg/mL **(Fig. 3a)**. Notably, the agglutination kinetics of all HM-IgGs were markedly faster and more complete than the parent IgG at each Ab concentrations and across all timepoints **(Fig. 3b)**.

We again assessed agglutination kinetics of FFIFF vs. IgG using whole semen. Similar to the PM sperm escape assay, substantially more FFIFF and the parent IgG were required to agglutinate sperm in native semen with comparable kinetics as washed sperm. FFIFF again exhibited markedly faster and more complete sperm agglutination kinetics than IgG at all mAb concentrations and all timepoints in whole semen **(Fig. 3d)**. Indeed, FFIFF agglutinated ≥90% of PM sperm within 30 s in 6 of 6 semen samples at 25 μg/mL, while IgG agglutinated ≥90% of PM sperm in 90 s in only one of six specimens at the same concentration **(Fig. 3c)**. Since lower sperm concentration (as found in semen from oligospermia, sub-fertile individuals) may limit agglutination potency due to reduced likelihood of sperm-sperm collision, and higher quantities of sperm may saturate the agglutination potential, we further assessed whether FFIFF can effectively reduce PM sperm at 1 million PM sperm/mL and 25 million PM sperm/mL. FFIFF maintained similar superior agglutination kinetics than IgG across both conditions **(Supplementary Fig. 3)**. These results underscore the increased potency for FFIFF compared to the parent IgG across diverse conditions.

### HM-IgGs preserve Fc-mucin crosslinking

Previous work has shown that IgG and IgM Abs can trap individual spermatozoa in mucus despite the continued vigorously beating action of the flagellum; clinically, this is referred to as the “shaking phenomenon”^9^. This muco-trapping function is similar to our recent observations with Herpes Simplex Virus (HSV) that multiple HSV-bound IgGs can form polyvalent adhesive interactions between their Fc domains and mucin fibers in CVM, resulting in effective trapping of individual viral particles in CVM and blocking vaginal transmission of HSV in mice^13^. Therefore, we next assessed whether the HM-IgGs can reduce progressive motility in the relatively thin (low viscosity) human cervicovaginal mucus (CVM). We fluorescently labeled human sperm and quantified their motion in CVM treated with different mAbs using multiple particle tracking^32^. All 3 constructs were able to reduce progressive motility of spermatozoa to the same extent as the parent IgG, indicating that addition of Fabs to both the N- and C-terminus of the IgG did not interfere with Fc-mucin crosslinking **(Fig. 4).**

**Fig. 4:**
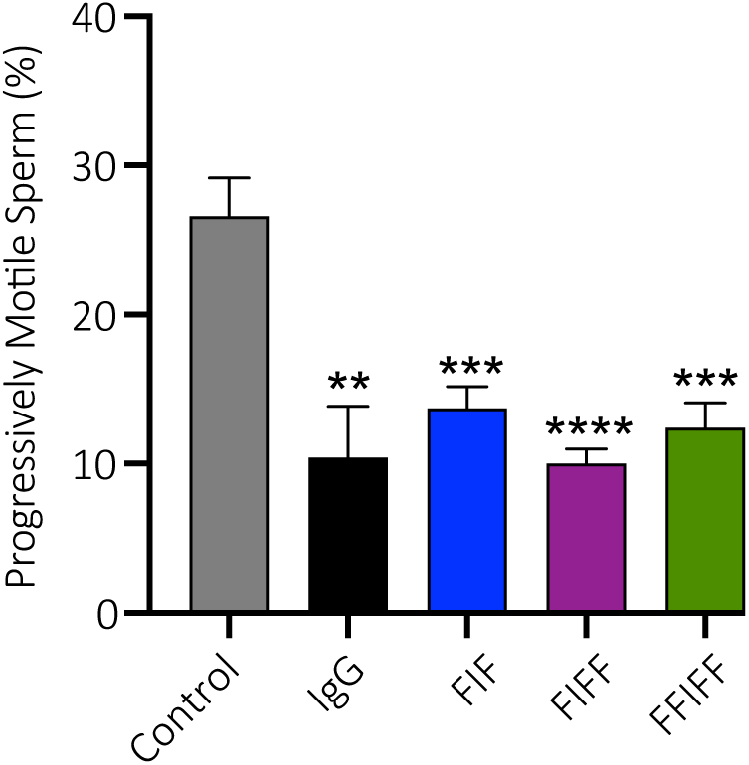
Highly multivalent anti-sperm IgG constructs conserve the trapping potency of the parent IgG. The trapping potency of the indicated Abs measured by quantifying the percentage of fluorescently labeled PM sperm in Ab-treated CVM using neural network tracker analysis software. 25 μg/mL of multivalent Abs or isotype control and purified motile sperm at the concentration of 5.8 x 10^4^ PM sperm/mL were used. Motavizumab (anti-RSV IgG) was used as the isotype control. Data were obtained from n = 6 independent experiments with 6 unique combinations of semen and CVM specimens. P values were calculated using a one-tailed t-test. *P<0.05, **P < 0.01, ***P < 0.001 and ****P < 0.0001. Lines indicate arithmetic mean values and standard error of mean.

### FIF and FFIFF effectively reduce PM sperm in sheep vagina

Since the unique glycoform of CD52g is only found in human and chimpanzee sperm^33^, there is no practical animal model to perform mating-based contraceptive efficacy studies. Instead, we designed a sheep study that parallels the human post-coital test (PCT)^34–38^, which assesses the reduction of PM sperm in the FRT given that PM sperm is required for fertilization. Clinical PCT studies have proven to be highly predictive of contraceptive efficacy in contraceptive effectiveness clinical trials^38–43^. The sheep vagina is physiologically and anatomically very similar to the human vagina^44,45^, making it the gold standard for assessing vaginal products. We instilled PBS, IgG, FIF or FFIFF into the sheep vagina, followed by brief simulated intercourse with a vaginal dilator (15 strokes), vaginal instillation of whole human semen, brief simulated intercourse (5 strokes), and finally, recovery of the semen mixture from the sheep vagina 2 mins post semen instillation for immediate assessment of sperm motility. The parent IgG, FIF and FFIFF all effectively reduced PM sperm at 333 μg dose **(Fig. 5).** At 10-fold lower concentrations i.e. 33 μg dose per sheep, FIF and FFIFF were still able to reduce PM sperm by 97% and >99%, respectively, whereas the parent IgG failed to substantially reduce PM sperm at the same concentration **(Fig. 5)**.

**Fig. 5:**
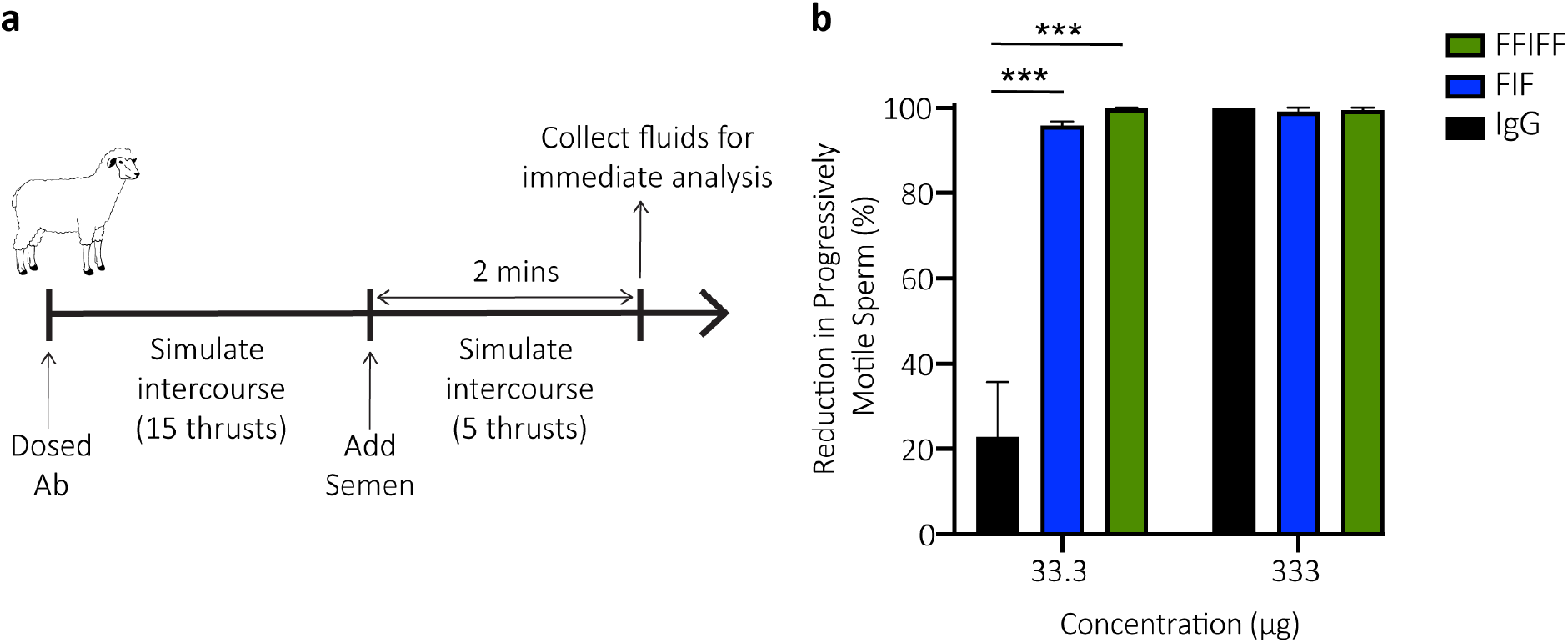
Highly multivalent anti-sperm IgG constructs demonstrate stronger agglutination potency than the parent IgG in surrogate sheep studies. (a) Schematic of the study design. (b) Potency of IgG, FIF and FFIFF measured by the reduction in percentage of PM sperm in sheep’s vaginal fluid after Ab-treatment compared to PBS-treatment. Data were obtained from n = 3 independent experiments. Treatment administration was blinded, and quantifications were manually performed in a blinded fashion. P values were calculated using a one-way ANOVA with Dunnett’s multiple comparisons test. *P < 0.05, **P < 0.01, and ***P < 0.001. Lines indicate arithmetic mean values and standard error of mean.

## Discussion

The exceptional potencies we observed in sheep despite the low total dose of mAb is a direct consequence of topical vaginal delivery. Since sperm is restricted to the FRT, topical delivery confines the contraceptive mAb directly at the site of action. Given the limited volume of secretions in the FRT (≤ ~1 mL in the human vagina^46^, relatively high concentration of mAb locally can be achieved even with very limited total dose of mAb. In contrast, systemically delivered mAbs must dilute into a large blood volume (~5L), distribution to non-target tissues, natural catabolic degradation, and limited distribution into the FRT, including the vagina. The markedly lower doses of mAb needed to sustain contraceptive levels in the FRT with vaginal delivery should translate to substantially less mAb needed, and consequently cost savings. Furthermore, by simply reinforcing the mucus barrier that is continuously secreted and cleared, rather than altering physiological mechanisms underpinning fertility (e.g. hormones), topical immunocontraception most likely affords rapid return to fertility, unlike the many months of delay experienced by some women even after they discontinued the use of long acting hormonal contraceptives.

Unlike small molecule contraceptives, contraceptive mAbs are likely to be exceptionally safe due to the specificity of targeting, particularly when targeted to unique epitopes present only on sperm and not on expressed in female tissues. Safety is likely to be further enhanced by topical delivery: mAb delivered to mucosal surfaces such as the vagina are poorly absorbed into the systemic circulation^47,48^, and the vagina represents a poor immunization inductive site, with limited immune response even when vaccinating with the aid of highly immunostimulatory adjuvants^49^. Finally, vaginal secretions already contain much higher levels of endogenous IgG (i.e. 1-2 mg/mL)^13^, making it unlikely that vaginal delivery of HM-IgGs comprised of fully human Fabs and Fc would trigger local toxicities.

Sperm must swim through mucus and ascend to the upper tract to reach and fertilize the egg. Typically, only ~1% of the ejaculated sperm enter the cervix, even fewer reaching the uterus, and only dozens of sperm (out of the ~200 million in the ejaculate) actually reach the neighborhood of the egg^50^. Accordingly, poor sperm motility in mid-cycle cervical mucus and low total sperm count are considered good correlates to low conception rates. Human semen averages between 45-65 million sperm/mL, 15 million sperm/mL marks the lowest 5th percentile in men with proven fertility, and <5 million sperm/mL is often considered severe oligozoospermia with very low fertility^29,30^. These observations suggest a marked reduction of progressive sperm motility, even if incomplete (e.g. 10-fold reduction in PM sperm fractions), would likely provide effective contraception. This expectation is also consistent with the observations that even under ideal circumstances, with unprotected intercourse on the cycle day of maximum fertility, the odds of conceiving is only about ~10%^51^. This indicates that only a small (i.e. limiting) number of motile sperm would reach the egg per intercourse; thus, reducing progressive sperm motility in the vagina and cervical canal should proportionally reduce likelihood of conceiving. These findings, together with the contraceptive success with topical ASA against rabbit sperm^22^, suggest arresting progressive sperm motility in mucus using mAb (which can reduce PM sperm by >99%) should provide an effective form of contraception.

Advances in vaginal drug delivery technologies have made available multiple methods for delivering contraceptive mAbs. For instance, a rapidly dissolving vaginal film^52^ could provide on-demand non-hormonal contraception. For a non-coitally associated method, HM-IgGs can be released from intravaginal rings (IVRs) that afford nearly constant, zero-order release kinetics of mAbs^53^. The fertility window in women typically starts a few days after the end of menses, and ends ~12-14 days before the start of the next menses^54,55^. Thus, for a 35-day menstrual cycle, an IVR would only need to release contraceptive mAbs for ~18 days if the IVR is inserted at the end of menses (~Day 5 of a typical cycle). A hypothetical 20 mg of FFIFF loaded per IVR could sustain >100 μg/mL^53^ concentrations in FRT secretions. With the cost of large scale mAb manufacturing at ~$100/g^56^ in 2009, and continued decrease in mAb manufacturing costs, we speculate the FFIFF Ab quantity required to provide monthlong non-hormonal contraception may costs no more than ~$2/month to produce.

Many of the current multivalent Abs are bispecific or trispecific in nature and must contend with potential mispairing of light and heavy chains. As a result, many such engineered Ab formats, such as single-chain variable fragment (scFv) or camel-derived nanobodies, involve substantial deviation from natural human Ab structure. scFv-based multivalent Ab constructs frequently suffer from low stability, heterogeneous expression, and decreased affinity and specificity stemming from the removal of the C_H_1/C_L_ interface present in a full-length Fab. The introduction of orthogonal mutations to facilitate heavy and light chain pairing can also substantially reduce mAb yield or overall stability. These limitations do not apply when generating monospecific HM-IgGs, which possess identical and full length human Fabs. Our strategy to covalently link additional Fabs to a parent heavy chain also contrasts with current multimerization strategies based on self-assembly of multiple IgGs based on Fc-mutations^57^ or appending an IgM tail-piece^58,59^, which often suffers poor homogeneity and stability. The combination of fully intact human Fabs and covalent linkages likely contributes to the surprising thermal stability, homogeneity and bioprocessing ease of the HM-IgGs developed here. We believe the IgG multimerization strategy presented here is likely a promising platform for developing mAbs where agglutination is a critical effector function.

## Methods

### Study design and ethics

*Ex vivo* studies with human semen as well as human cervicovaginal mucus (CVM) utilized specimens obtained following a protocol approved by the Institutional Review Board (IRB) of the University of North Carolina at Chapel Hill (IRB-101817). Informed written consent was obtained from all male and female subjects before the collection of any material. Subjects were recruited from the Chapel Hill and Carrboro, NC area in response to mass student emails and ads in print media. For the sheep surrogate post-coital test performed at the University of Texas Medical Branch (UTMB), UTMB IRB approval and written consent allowed for collection of fresh semen from pre-screened male volunteers on the day of each sheep study. Sheep studies were approved by the UTMB Institutional Animal Care and Use Committee (IACUC) and utilized 5 female Merino crossbred sheep.

### Construction of the parent IgG and HM-IgGs plasmids

The variable heavy (V_H_) and variable light (V_L_) DNA sequences for anti-sperm IgG1 antibody (Ab) were obtained from the published sequence of H6-3C4 mAb^26^.

For the construction of expression plasmid encoding the light chain (LC), the gene fragment consisting of V_L_ and C_L_ (C_λ_) DNA sequences was synthesized using a custom gene-synthesis service (Integrated DNA Technologies) and cloned in-house into a customized empty mammalian expression vector (pAH, ThermoFisher Scientific) using KpnI (5’) and EcoRI (3’) restriction sites.

For the construction of expression plasmids encoding heavy chains (HC) for parent and HM-IgG Abs, four cloning vectors comprising of V_H_/C_H_1-(G_4_S)_6_ Linker-V_H_, V_H_/C_H_1-(G_4_S)_6_ Linker-V_H_/C_H_1-(G_4_S)_6_ Linker-V_H_, (G_4_S)_6_ Linker-V_H_/C_H_1 and (G_4_S)_6_ Linker-V_H_/C_H_1-(G_4_S)_6_ Linker-V_H_/C_H_1 sequences were synthesized using GeneArt^®^ gene synthesis service (ThermoFisher Scientific).

For the construction of expression plasmid encoding HC for IgG, V_H_ fragment was amplified from the cloning vector comprising of V_H_/C_H_1-(G_4_S)_6_ Linker-V_H_ using forward primer, 5’ TAAGCAGGTACCGCCACCATGAAGTG 3’, and reverse primer, 5’ TGCTTAGCTAGCTGGAGAAACTGTC 3’, and then cloned into a mammalian IgG1 expression vector (pAH comprising C_H_1-C_H_2-C_H_3 DNA sequence) using KpnI (5’) and NheI (3’) restriction sites.

For the construction of expression plasmid encoding HC for FIF, V_H_/C_H_1-(G_4_S)_6_ Linker-V_H_ fragment was first cloned into a mammalian IgG1 expression vector using KpnI (5’) and NheI (3’) restriction sites followed by the cloning of (G4S)6 Linker-VH/CH1 fragment using BamHI (5’) and MluI (3’) restriction sites.

For the construction of expression plasmid encoding HC for FIFF, V_H_/C_H_1-(G_4_S)_6_ Linker-V_H_ fragment was first cloned into the mammalian IgG1 expression vector using KpnI (5’) and NheI (3’) restriction sites followed by the cloning of (G_4_S)_6_ Linker-V_H_C_H_1-(G_4_S)_6_ Linker-V_H_/C_H_1 fragment using BamHI (5’) and MluI (3’) restriction sites.

For the construction of expression plasmid encoding HC for FFIFF, V_H_/C_H_1-(G_4_S)_6_ Linker-V_H_/C_H_1-(G_4_S)_6_ Linker-V_H_ fragment was first cloned into the same mammalian expression vector using KpnI (5’) and NheI (3’) restriction sites followed by the cloning of (G_4_S)_6_ Linker-V_H_/C_H_1-(G_4_S)_6_ Linker-V_H_/C_H_1 using BamHI (5’) and MluI (3’) restriction sites.

For the ligation of all HCs as well as the LC into mammalian expression vectors, a quick ligation kit (New England Biolabs) was used. All ligated DNA constructs were chemically transformed into TOP10 *E. coli* cells (Life Technologies) and plated on ampicillin plates for selection. Several bacterial colonies were picked from each plate and cultured, followed by plasmid miniprep (Qiagen MiniPrep Kit). Correct assembly of the HC and LC sequences into expression vectors were confirmed by sanger sequencing of the purified plasmids (Eurofins Genomics).

### Expression and purification of the parent IgG and HM-IgGs

The sequencing-confirmed expression plasmids encoding the HC and LC sequences of parent and HM-IgG Abs were chemically transformed and cultured in a 100 mL Terrific broth media overnight. Midiprep plasmid purifications were done using NucleoBond^®^ Xtra Midi EF Kits (Macherey-Nagel), according to the manufacturer’s protocols. Purified HC and LC plasmids were transfected into Expi293F cells using ExpiFectamine™ 293 Transfection reagents, according to the manufacturer’s protocols (Gibco). For IgG, HC and LC plasmid were co-transfected using a 1:1 ratio at 1 μg total DNA per 1 mL of culture. For FIF, HC and LC plasmid were co-transfected using a 1:3 ratio at 1 μg total DNA per 1 mL culture. For FIFF, HC and LC plasmid were co-transfected using a 1:4 ratio at 1 μg total DNA per 1 mL culture. For FFIFF, HC and LC plasmid were co-transfected using a 1:5 ratio at 1 μg total DNA per 1 mL culture. Transfected Expi293F cells were grown at 37°C in a 5% CO_2_ incubator and shaken at 125 r.p.m. for 3-5 days. Supernatants were harvested by centrifugation at 12,800 g for 10 min, passed through 0.22 μm filters and purified using standard protein A/G chromatography method. Purified Abs were quantified using absorbance at 280 nm along with corresponding protein extinction coefficients.

### Antibody characterization using SDS-PAGE, SEC-MALS and nano-DSF

SDS-PAGE experiments were performed using 4-12% NuPage Bis-Tris gels (Invitrogen) in 1x NuPage MOPS buffer under both reducing and non-reducing conditions to confirm the molecular weight and assembly of all Ab constructs. For each sample, 1 μg of protein was diluted in 3.75 μL LDS sample buffer followed by the addition of 11.25 μL nuclease-free water. Proteins were then denatured at 70°C for 10 min. Next, 0.3 μL of 0.5 M tris (2-carboxyethyl) phosphine (TCEP) was added as a reducing agent to the denatured protein for reduced samples and incubated at room temperature (RT) for 5 min. Bio-Rad Precision Protein Plus Unstained Standard and Novex™ Sharp Pre-stained Protein Standard were used as ladders. After loading the samples, the gel was run for 50 min at a constant voltage of 200 V. The protein bands were visualized by staining with Imperial Protein Stain (Thermo Scientific) for 1 hr followed by overnight de-staining with Milli-Q water. Image J software (Fiji) was used to adjust the brightness and contrasts of the SDS-PAGE gel for visual purposes.

SEC-MALS experiments were performed at RT using a GE Superdex 200 10/300 column connected to an Agilent FPLC system, a Wyatt DAWN HELEOS II multi-angle light-scattering instrument, and a Wyatt T-rEX refractometer. The flow rate was maintained at 0.5 mL/min. The column was equilibrated with 1X PBS, pH 7.4 containing 200 mg/L of NaN_3_ before sample loading. 50-100 μg of each sample was injected onto the column, and data were collected for 50 min. The MALS data were collected and analyzed using Wyatt ASTRA software (Ver. 6).

NanoDSF experiments were performed using a Nanotemper Prometheus NT.48 system. Samples were diluted to 0.5 mg/mL in 1X PBS at pH 7.4 and loaded into Prometheus NT.48 capillaries. Thermal denaturation experiments were performed from 25°C to 95°C at the rate of 1°C/min, measuring the intrinsic tryptophan fluorescence at 330 nm and 350 nm. The melting temperature for each experiment was calculated automatically by Nanotemper PR. Thermcontrol software by plotting the ratiometric measurement of the fluorescent signal against increasing temperature. The aggregation temperature for each experiment was also calculated automatically by Nanotemper PR. Thermcontrol software via the detection of the back-reflection intensity of a light beam that passes the sample.

### Semen collection and isolation of motile sperm

Healthy male subjects were asked to refrain from sexual activity for at least 24 hr prior to semen collection. Semen was collected by masturbation into sterile 50 mL sample cups and incubated for a minimum of 15 min post-ejaculation at RT to allow liquefaction. Semen volume was measured, and the density gradient sperm separation procedure (Irvine Scientific) was used to extract motile sperm from liquefied ejaculates. Briefly, 1.5 mL of liquified semen was carefully layered over 1.5 mL of Isolate^®^ (90% density gradient medium, Irvine Scientific) at RT, and centrifuged at 300 g for 20 min. Following centrifugation, the upper layer containing dead cells and seminal plasma was carefully removed without disturbing the motile sperm pellet in the lower layer. The sperm pellet was then washed twice with the sperm washing medium (Irvine Scientific) by centrifugation at 300 g for 10 min.

Finally, the purified motile sperm pellet was resuspended in the sperm washing medium, and an aliquot was taken for determination of sperm count and motility using computer-assisted sperm analysis (CASA). All semen samples used in the functional assays exceeded lower reference limits for sperm count (15 × 10^6^ total sperm/mL) and total motility (40%) as indicated by WHO guidelines^29^.

### Assessment of CD52g expression in human semen samples

The expression of the CD52g target antigen on the sperm of human semen donors was assessed on de-identified residual aliquots of semen samples from 100 donors presenting to an andrology clinic for semen quality evaluation (see Supplementary Table 2 for ethnic background). Aliquots of semen were placed on glass slides and an equal volume of H6-3C4 IgM mAb directed against the CD52g antigen (tissue culture supernatant, gift of S. Isojima^10^), was added and mixed briefly with a pipette tip. A coverslip was applied, and agglutination was assessed microscopically 3 minutes after Ab addition via a 20X objective. Strong agglutination was observed in all samples. The study protocol was approved by the Institutional Review Board of the Union Memorial Hospital, Baltimore, MD.

### Sperm count and motility using CASA

The Hamilton-Thorne computer-assisted sperm analyzer, 12.3 version, was used for the sperm count and motility analysis in all experiments unless stated otherwise. This device consists of a phasecontrast microscope (Olympus CX41), a camera, an image digitizer and a computer with a Hamilton-Thorne Ceros 12.3 software to save and analyze the acquired data. For each analysis, 4.4 μL of the semen sample was inserted into MicroTool counting chamber slides (Cytonix). Then, six randomly selected microscopic fields, near the center of the slide, were imaged and analyzed for progressively motile (PM) and non-progressively motile (NPM) sperm count. The parameters that were assessed by CASA for motility analysis were as follows: average pathway velocity (VAP: the average velocity of a smoothed cell path in μm/s), the straight-line velocity (VSL: the average velocity measured in a straight line from the beginning to the end of track in μm/s), the curvilinear velocity (VCL: the average velocity measured over the actual point-to-point track of the cell in μm/s), the lateral head amplitude (ALH: amplitude of lateral head displacement in μm), the beat cross-frequency (BCF: frequency of sperm head crossing the sperm average path in Hz), the straightness (STR: the average value of the ratio VSL/VAP in %), and the linearity (LIN: the average value of the ratio VSL/VCL in %). PM sperm were defined as having a minimum of 25 μm/s VAP and 80% of STR^27^. The complete parameters of the Hamilton-Thorne Ceros 12.3 software are listed in Supplementary Table 2.

### Whole sperm ELISA

Briefly, half-area polystyrene plates (CLS3690, Corning) were coated with 2 x 10^5^ sperm per well in 50 μL of NaHCO_3_ buffer (pH 9.6). After overnight incubation at 4°C, the plates were centrifuged at the speed of 300 g for 20 min. The supernatant was discarded, and the plates were air-dried for 1 hr at 45°C. The plates were washed once with 1X PBS. 100 μL of 5% milk was incubated at RT for 1 hr to prevent non-specific binding of Abs to the microwells. The serial dilution of mAbs in 1% milk was added to the microwells and incubated overnight at 4°C. Motavizumab, a mAb against the respiratory syncytial virus, was constructed and expressed in the laboratory by accessing the published sequence and used as a negative control for this assay^60^. After primary incubation, the plates were washed three times using 1X PBS. Then, the secondary Ab, goat anti-human IgG F(ab’)_2_ Ab HRP-conjugated (1:10,000 dilutions in 1% milk, 209-1304, Rockland Inc.) was added to the wells and incubated for 1 hr at RT. The washing procedure was repeated and 50 μL of the buffer containing substrate (1-Step Ultra TMB ELISA Substrate, Thermo Scientific) was added to develop the colorimetric reaction for 15 min. The reaction was quenched using 50 μL of 2N H_2_SO_4_, and the absorbance at 450 nm (signal) and 570 nm (background) was measured using SpectraMax M2 Microplate Reader (Molecular Devices). Each experiment was done with samples in triplicates and repeated two times as a measure of assay variability.

### Scanning electron microscopy

Briefly, 20 x 10^6^ washed sperm was centrifuged at 300 g for 10 min. The supernatant was discarded without disturbing the sperm pellet. Next, 200 μl of anti-sperm IgG constructs or 1X PBS was added to the sperm pellet. The Ab-sperm solution was mixed by pipetting and incubated for 5 mins using an end-over-end rotator. 200 μl of 4% PFA prepared in 0.15 M Sodium Phosphate buffer was added to the Ab-sperm solution and incubated for 10 min using an end-over-end rotator. 50 μl of fixed sperm samples was filtered through membrane filters (10562, K04CP02500, Osmonics) with 5 mL of 0.15 M Sodium Phosphate buffer. The filter was washed one more time with 0.15 M Sodium Phosphate buffer. Next, the samples were dehydrated in a graded series of alcohol (30% ethanol, 50% ethanol, 70% ethanol, 90% ethanol, 100% ethanol x 3) for 10 minutes each. Filters were transferred to a plate with the transitional solvent, Hexamethyldisilazane (Electron Microscopy Sciences) and allowed to dry after one exchange. Filters were adhered to aluminum stubs with carbon adhesives and samples were sputter-coated with gold-palladium alloy (Au:Pd 60:40 ratio, 91112, Ted Pella Inc.) to a thickness of 3 nm using Cressington Sputter Coater 208 hr. Six random images were acquired for each sample using a Zeiss Supra 25 FESEM with an SE2 Electron detector at 2500X magnification.

### Sperm escape assay

This assay was conducted using whole semen and purified motile sperm at the final concentration of 5 x 10^6^ PM sperm/mL. Briefly, 40 μL aliquots of purified motile sperm or whole semen were transferred to individual 0.2 mL PCR tubes. Sperm count and motility were performed again on each 40 μL aliquot using CASA. This count serves as the original (untreated) concentration of sperm for evaluating the agglutination potencies of respective Ab constructs. Following CASA, 30 μL of purified motile sperm or native semen was added to 0.2 mL PCR tubes containing 30 μL of Ab constructs, and gently mixed by pipetting. The tubes were then held fixed at 45° angles in a custom 3D printed tube holder for 5 min at RT. Following this incubation period, 4.4 μL was pipetted from the top layer of the mixture with minimal perturbation of the tube and transferred to the CASA instrument to quantify the number of PM sperm. The percentage of the PM sperm that escaped agglutination was computed by dividing the sperm count obtained after treatment with Ab constructs by the original untreated sperm count in each respective tube, correcting for the 2-fold dilution with Ab. Each experimental condition was evaluated in duplicates on each semen specimen, and the average from the two experiments was used in the analysis. At least 6 independent experiments were done with at least n=4 unique semen samples.

### Agglutination kinetics assay

This assay was conducted using both whole semen and purified motile sperm at the final concentration of 1 x 10^6^ PM sperm/mL, 5 x 10^6^ PM sperm/mL and 25 x 10^6^ PM sperm/mL. Briefly, 4.4 μL of purified motile sperm or whole semen was added to 4.4 μL of Ab constructs in 0.2 mL PCR tubes, and mixed by gently pipetting up and down three times over 3 s. A timer was started immediately while 4.4 μL of the mixture was transferred to chamber slides with a depth of 20 μm (Cytonix), and video microscopy (Olympus CKX41) using a 10X objective lens focused on the center of chamber slide was captured up to 90 s at 60 frames/s. PM sperm count was measured by CASA every 30 s up to 90 s. The reduction in percentage of the PM sperm at each time point was computed by normalizing the PM sperm count obtained after Ab-treatment to the PM sperm count obtained after treatment with sperm washing medium. Each experimental condition, except for 25 x 10^6^ PM sperm/mL, was evaluated in duplicates on each semen specimen, and the average from the two experiments was used in the analysis. At least 6 independent experiments were done with at least n=4 unique semen samples.

### Cervicovaginal mucus collection and processing

Cervicovaginal mucus (CVM) was collected as previously described^13^. Briefly, undiluted CVM secretions, averaging 0.5 g per sample, were obtained from women of reproductive age, ranging from 20 to 44 years old, by using a self-sampling menstrual collection device (Instead Softcup). Participants inserted the device into the vagina for at least 30 s, removed it, and placed it into a 50 mL centrifuge tube. Samples were centrifuged at 230 g for 5 min to collect the secretions. Samples were collected at various times throughout the menstrual cycle, and the cycle phase was estimated based on the last menstrual period date normalized to a 28-day cycle. Samples that were non-uniform in color or consistency were discarded. Donors stated they had not used vaginal products nor participated in unprotected intercourse within 3 days before donating. All samples had pH < 4.5.

### Fluorescent labeling of sperm

Purified motile sperm were fluorescently labeled using Live/Dead Sperm Viability Kit (Invitrogen Molecular Probes), which stains live sperm with SYBR 14 dye, a membrane-permeant nucleic acid stain, and dead sperm with propidium iodide (PI), a membrane impermeant nucleic acid stain. Briefly, SYBR 14 stock solution was diluted 50-fold in sperm washing media. Next, 5 μL of diluted SYBR 14 and PI dye were added to 1 mL of washed sperm resulting in final SYBR 14 and PI concentration of 200 nM and 12 μM respectively. The sperm-dye solution was incubated for 10 min at 36°C. Then, the solution was washed twice using the sperm washing medium to remove unbound fluorophores by centrifuging at 300 g for 10 min. Next, the labeled motile sperm pellet was resuspended in the sperm washing medium, and an aliquot was taken for determination of sperm count and motility using CASA.

### Multiple particle tracking studies

To mimic the dilution and neutralization of CVM by alkaline seminal fluid, CVM was first diluted threefold using sperm washing medium and titrated to pH 6.8-7.1 using small volumes of 3 N NaOH. The pH was confirmed using pH test strips. Next, 4 μL of Ab constructs or control (anti-RSV IgG1) was added to 60 μL of diluted and pH adjusted CVM and mixed well in a CultureWell™ chamber slides (Invitrogen) followed by addition of 4 μL of 1 x 10^6^ PM sperm/mL of fluorescently labeled sperm. Once mixed, sperm, Ab, and CVM were incubated for 5 min at RT. Then, translational motions of the sperm were recorded using an electron-multiplying charge-coupled-device camera (Evolve 512; Photometrics, Tucson, AZ) mounted on an inverted epifluorescence microscope (AxioObserver D1; Zeiss) equipped with an Alpha Plan-Apo 20/0.4 objective, environmental (temperature and CO_2_) control chamber, and light-emitting diode (LED) light source (Lumencor Light Engine DAPI/GFP/543/623/690)^13^. 15 videos (512 × 512 pixels, 16-bit image depth) were captured for each Ab condition with MetaMorph imaging software (Molecular Devices) at a temporal resolution of 66.7 ms and spatial resolution of 50 nm (nominal pixel resolution, 0.78 μm/pixel) for 10 s. Next, the acquired videos were analyzed via a neural network tracking software modified with standard sperm motility parameters to determine the percentage of PM sperm^32^. At least 6 independent experiments were performed, each using a unique combination of CVM and semen specimens.

### In vivo surrogate efficacy studies

On the test day, each sheep received a randomized unique Ab treatment and all sheep were dosed with the same semen mixture that was pooled from 3-5 donors. Briefly, 1 mL of anti-sperm Abs or PBS control, provided under blind to the animal facility, were instilled into sheep’s vagina and thoroughly mixed using a vaginal dilator for 15 strokes. Next, 1 mL of pooled whole semen was pipetted into sheep’s vagina, followed by a simulated intercourse with vaginal dilator for 5 strokes. Two minutes after introduction of semen, fluids from the sheep vagina were recovered and assessed for the PM sperm count in a hemocytometer (Bright-Line™ Hemacytometer) under light microscope (Olympus IX71) using a 20X objective with Thorlabs camera. Each Ab condition was repeated two more times in the same group of sheep (n=5) with at least 7 days interval in between experiments. Treatments and quantifications were performed in a blinded fashion.

### Statistical analysis

All analyses were performed using GraphPad Prism 8 software. For multiple group comparisons, P values were calculated using a one-way ANOVA with Dunnett’s multiple comparisons tests. To compare the percent reduction of PM sperm by IgG vs FFIFF, using whole semen as well as washed sperm at the concentration of 1 x 10^6^ PM sperm/mL and 25 x 10^6^ PM sperm/mL, one-tailed t-test was performed. Similarly, the comparison between control- and anti-sperm Ab-treated fluorescent PM sperm was performed using a one-tailed t-test. In all analyses, α=0.05 for statistical significance. All data are presented as the mean ± standard error of the mean.

## Acknowledgements

We thank Dr. Deborah O’Brien for providing the CASA instrument and her assistance in setting up the CASA measurements. Special thanks to the UNC Macromolecular Interactions Facility for instruments used in SEC-MALS and DSF studies, and Microscopy Services Laboratory (MSL) for instruments used in SEM experiments. This work was financially supported by the Eshelman Institute of Innovation (S.K.L.); The David and Lucile Packard Foundation (2013-39274; S.K.L); National Institutes of Health under grants R56HD095629 (S.K.L.), U54HD096957 (T.R.M. and S.K.L.), R43HD094454 (T.R.M.) and R44HD097063 (T.R.M.); National Science Foundation (DMR-1810168; S.K.L.); and PhRMA Foundation Graduate Fellowship (B.S.).

## Author contributions

B.S, S.K.L and T.R.M conceptualized the study. B.S and S.K.L designed experiments and wrote the manuscript. K.V and T.R.M edited the manuscript. B.S, T.M.J and A.S engineered and assembled the DNA constructs for multivalent Abs. B.S planned and carried out the expression, purification, and characterization of the multivalent Abs. B.S and E.C.C planned and performed sperm escape and agglutination kinetics assays using purified sperm at the concentration of 5 x 10^6^ PM sperm/mL. B.S planned and carried out the remaining sperm escape and agglutination kinetics assays and SEM. B.S planned and conducted sperm trapping assay, and A.S developed an algorithm and analyzed the trapping data. S.S.O performed the assessment of CD52g expression frequency in human semen samples. B.S, S.K.L, T.R.M and K.V designed the experiments for sheep studies. J.S, Z.Y and K.V planned and conducted sheep studies, followed by data analysis by B.S, J.S, Z.Y and K.V.

## Data availability

All data are available from the corresponding author upon reasonable request.

## Code availability

Code used to analyze fluorescent PM sperm in vaginal mucus samples is available from the corresponding author upon reasonable request.

