## Supplementary File for "Engineering highly multivalent sperm-binding IgG antibodies for potent non-hormonal female contraception"

**Supplementary Information**

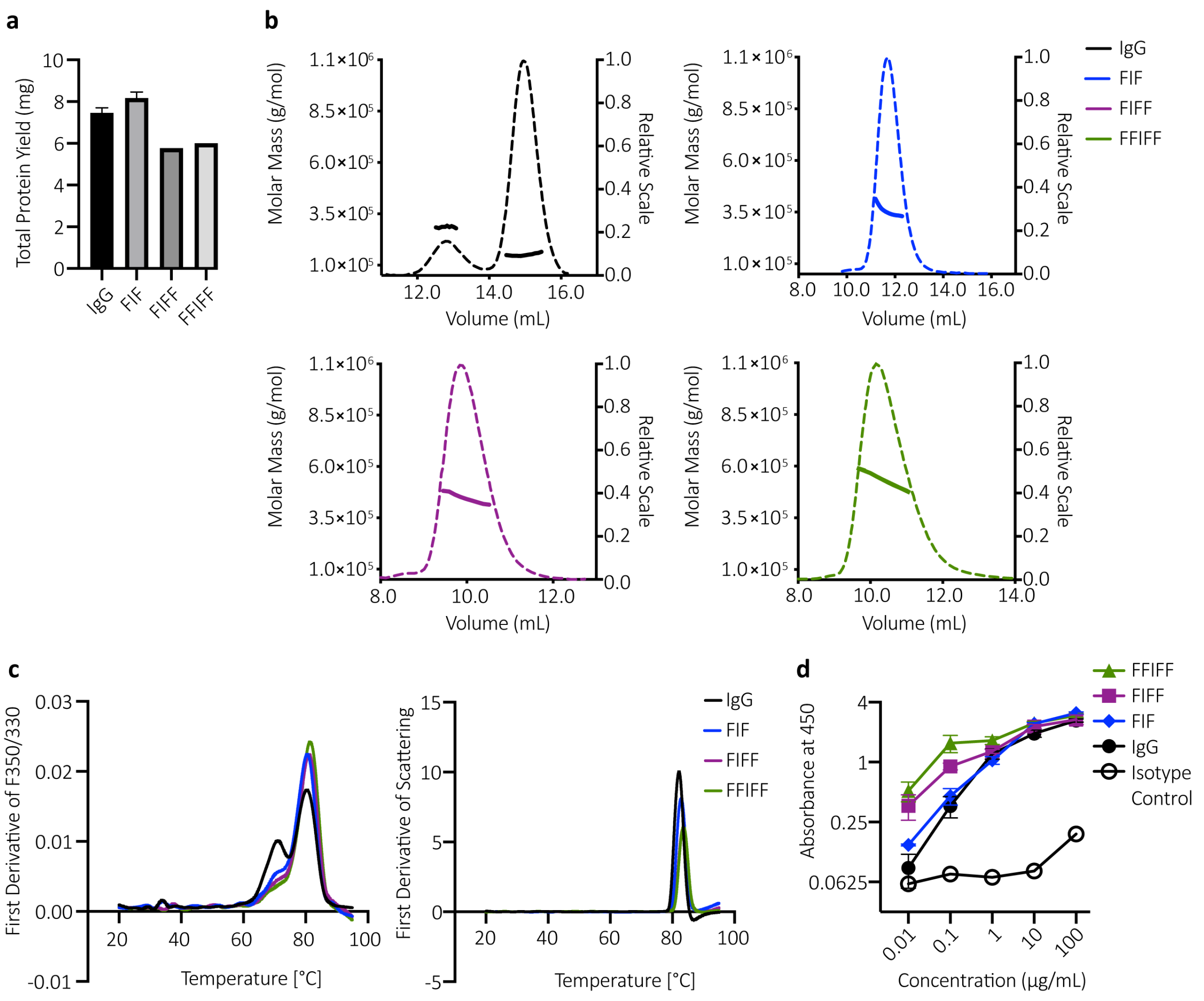


**Supplementary Fig. 1: Additional characterization of highly multivalent anti-sperm IgG antibodies.**

(a) Production yield of IgG, FIF, FIFF and FFIFF, purified from 90 mL Expi293 transfection. Data were obtained from n = 2 independent transfection for IgG and FIF, and one experiment for FIFF and FFIFF. (b) SEC-MALS curves of the parent IgG, FIF, FIFF and FFIFF. Thick lines indicate the calculated molecular mass (left y-axis) and the dotted lines show the homogenous profile (right y-axis) of each antibody. Data collected from one experiment. (c) The melting temperatures (left) and aggregating temperature (right) of the indicated antibodies as determined by nanoDSF by measuring intrinsic fluorescence and changes in back-reflection of proteins respectively. The experiment was performed in duplicates and averaged. (d) Whole sperm ELISA to assess the binding potency of the indicated antibodies to human sperm. Motavizumab (anti-RSV IgG) was used as the isotype control. Data were obtained from n = 3 independent experiments with n = 3 unique semen donors. Each experiment was performed in triplicates and averaged. Lines indicate arithmetic mean values and standard error of mean.

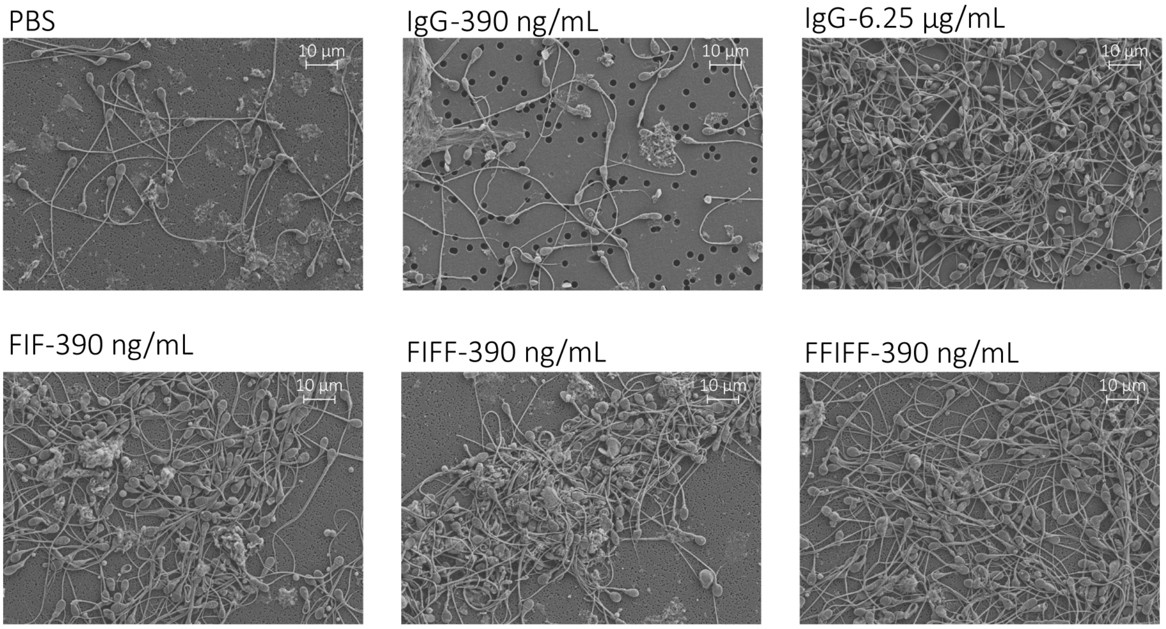


**Supplementary Fig. 2: Scanning electron microscopy images of the washed sperm treated with anti-sperm IgGs for 5 min.** Images were obtained at 2500x magnification. Scale bar, 10 μm.

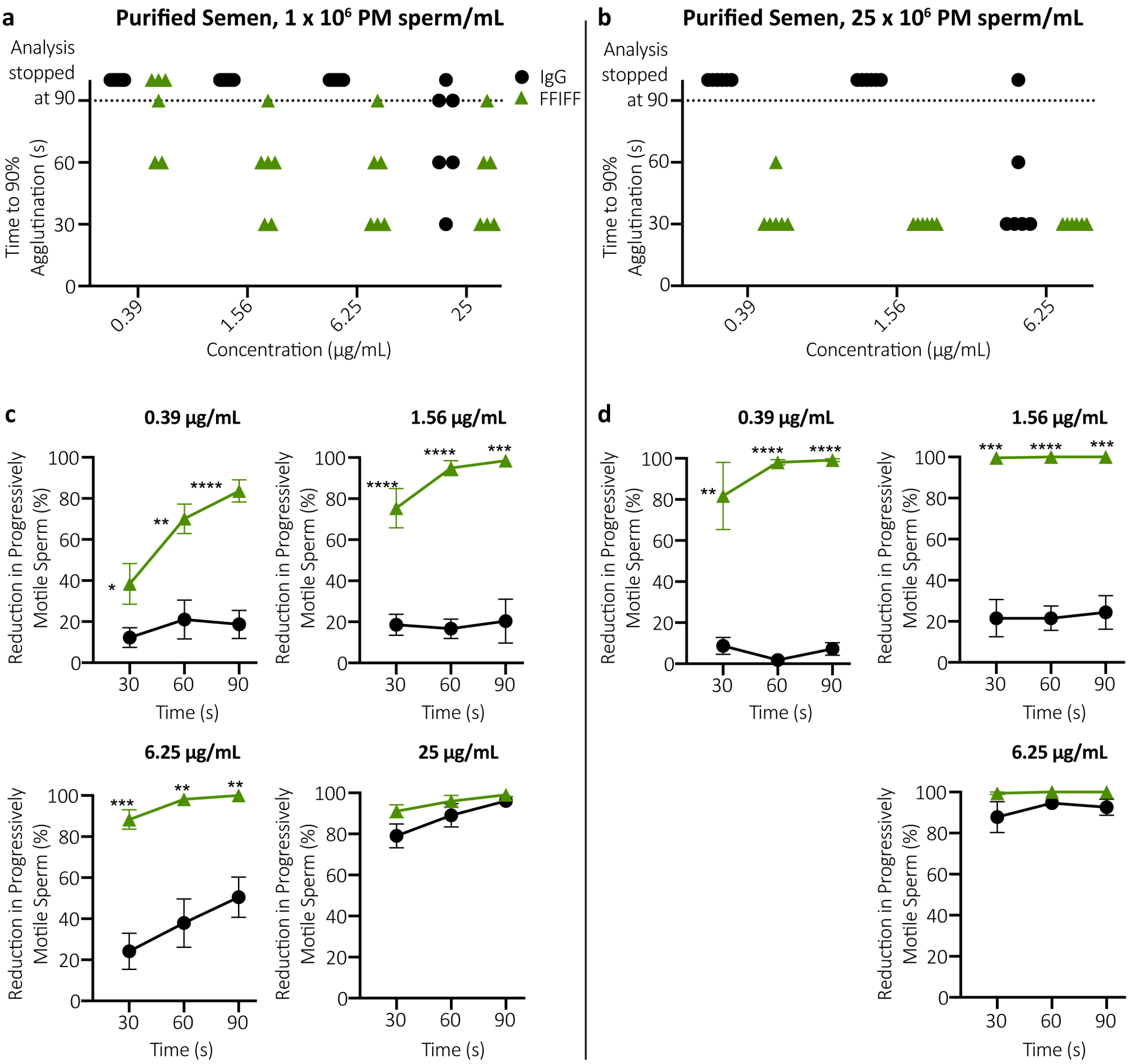


**Supplementary Fig. 3: FFIFF demonstrates faster agglutination kinetics than the parent IgG at both low and high sperm concentration.**

(a) Sperm agglutination kinetics of the parent IgG and FFIFF measured by the quantification of time required to achieve 90% agglutination of PM sperm compared to sperm washing media control using 1 x $10^6$ PM sperm/mL and (b) 25 x $10^6$ PM sperm/mL. (c) The rate of sperm agglutination for the parent IgG and FFIFF measured by the reduction in percentage of PM sperm at three different timepoints after Ab-treatment compared to sperm washing media control using 1 x $10^6$ PM sperm/mL and (d) 25 x $10^6$ PM sperm/mL. Data were obtained from n = 6 independent experiments with n = 6 different semen donors. Experiment involving 1 x $10^6$ PM sperm/mL was performed in duplicates and averaged. P values were calculated using a one-tailed t-test. *$P < 0.05$, **$P < 0.01$, ***$P < 0.001$ and ****$P < 0.0001$. Lines indicate arithmetic mean values and standard error of mean.

**Supplementary Table 1. The sperm motility parameters of the Hamilton-Thorne Ceros 12.3.**

| Parameter | Value | Parameter | Value |
|---|---|---|---|
| Frames Per Sec | 60 | Path Velocity (VAP) | 25 μm/s |
| No. of Frames | 60 | Straightness (STR) | 80 % |
| Minimum Cell Size | 3 pixels | VAP Cutoff | 10 μm/s |
| Default Cell Size | 6 pixels | VSL Cutoff | 0 μm/s |
| Minimum Contrast | 80 | Slow Cells | Motile |
| Default Cell Intensity | 20 | Standard Objective | 10x |
| Chamber Depth | 20 μm | Magnification | 1.87 |

**Supplementary Table 2. The demographics of the 100 semen samples agglutinated with parent IgG.**

| Race | Composition (%) |
|---|---|
| Caucasian | 73 |
| African-American | 26 |
| Asian | 1 |
